# Correlation analysis among single nucleotide polymorphisms in thirteen language genes and culture/education parameters from twenty-six countries

**DOI:** 10.1101/2021.08.22.457292

**Authors:** Bo Sun, Changlu Guo, Zhizhou Zhang

## Abstract

Language is a vital feature of any human culture, but whether language gene polymorphisms have meaningful correlations with some cultural characteristics during the long-run evolution of human languages largely remains obscure (uninvestigated). This study would be an endeavor example to find evidences for the above question’s answer. In this study, the collected basic data include 13 language genes and their randomly selected 111 single nucleotide polymorphisms (SNPs), SNP profiles, 29 culture/education parameters, and estimated cultural context values for 26 representative countries. In order to undertake principal component analysis (PCA) for correlation search, SNP genotypes, cultural context and all other culture/education parameters have to be quantitatively represented into numerical values. Based on the above conditions, this study obtained its preliminary results, the main points of which contain: (1) The 111 SNPs contain several clusters of correlational groups with positive and negative correlations with each other; (2) Low cultural context level significantly influences the correlational patterns among 111 SNPs in the principal component analysis diagram; and (3) Among 29 culture/education parameters, several basic characteristics of a language (the numbers of alphabet, vowel, consonant and dialect) demonstrate least correlations with 111 SNPs of 13 language genes.

## Introduction

Numerous studies are undertaken on food and human living environment, especially biological and physiological consequences they bring to human body, including effects on gene expression in different tissues. Meanwhile, much less studies are taken on similar effects resulted from human cultural and educational activities (CEAs). CEAs do not directly bring in and out of any eye-visible substance from human body (as food and human living environment do), but only induce psychological consequences plus neurological recognition-like output. In fact, more and more evidences have been gathered in that ECAs are capable of reshaping gene expression profiles of human body^[1–6]^; ECAs-induced elevation of brain recognition level will influence people’s daily life and directly bring changes in behaviors for food intake and many other human activities, thus indirectly bring in and out of eye-visible substance from human body. In this way, CEAs shall be associated with human gene evolution and revolution. Indeed, ECAs still belong to human phenotypic behavior, while any phenotype has innate interaction with genotype and influences each other in a long run.

Language genes are a group of specific genes in charge of unique human language functions. In the past twenty years, some progress has been seen on language gene polymorphisms from different ethnics^[7–12]^. These polymorphisms are primarily derived from long-term interactions between human and surrounding geographical environments to get adapted. Most of these polymorphisms are even likely produced and optimized in the early stage of human emergence. In a specific profile of geographical conditions, there is normally some type of ECAs for a certain human community that gets adapted to the specific geographical conditions. To what extent these ECAs can participate in the evolution of language gene polymorphism, is an interesting and not yet investigated a question.

As geographical factors, cultural and educational parameters also occur and evolve in a large scale space (at least in a scale as small as a village), and are hard to be modeled in a laboratory way in which variables can be changed one by one to observe relevant consequences in other variables. Even if we can employ a small number of people in a village-like space to perform culture-education activity-based research, it is still a big challenge to keep the observation for enough long a time (such as 20-50 years). As we know, human gene mutation happens all the time, but a stable polymorphism point in a genome is through selection in many generations before it can stably exist in a population. This means heritable SNPs with a stable pattern are normally accumulated in a long time as several hundred or thousand years, even longer. So what we can do is to employ correlation analysis method(s) to tackle questions as to what extent culture/education parameters play a role in reshaping the profile of language gene polymorphisms.

This study focused on correlation analysis among a total 111 SNPs from these 13 genes and a series of culture/education factors collected from 26 countries, and the author did find some interesting correlational parameters, including several strongest positive and negative correlations.

## Methods

### Language genes and their SNPs

Language is an emergent complicated function of human being, though many other animals also have their own ‘languages’. If a gene mutation is statistically or experimentally associated with a certain language function loss, it would be called language gene. Language gene SNP data were all randomly selected for each gene in the dbSNP database: https://www.ncbi.nlm.nih.gov/snp/; Table 1 listed 13 language genes, and a total 111 SNPs from these 13 genes were randomly selected for this study (Table 2, Supplementary file-1). Quantification of SNPs was described in Supplementary file-2.

**Table 1.**
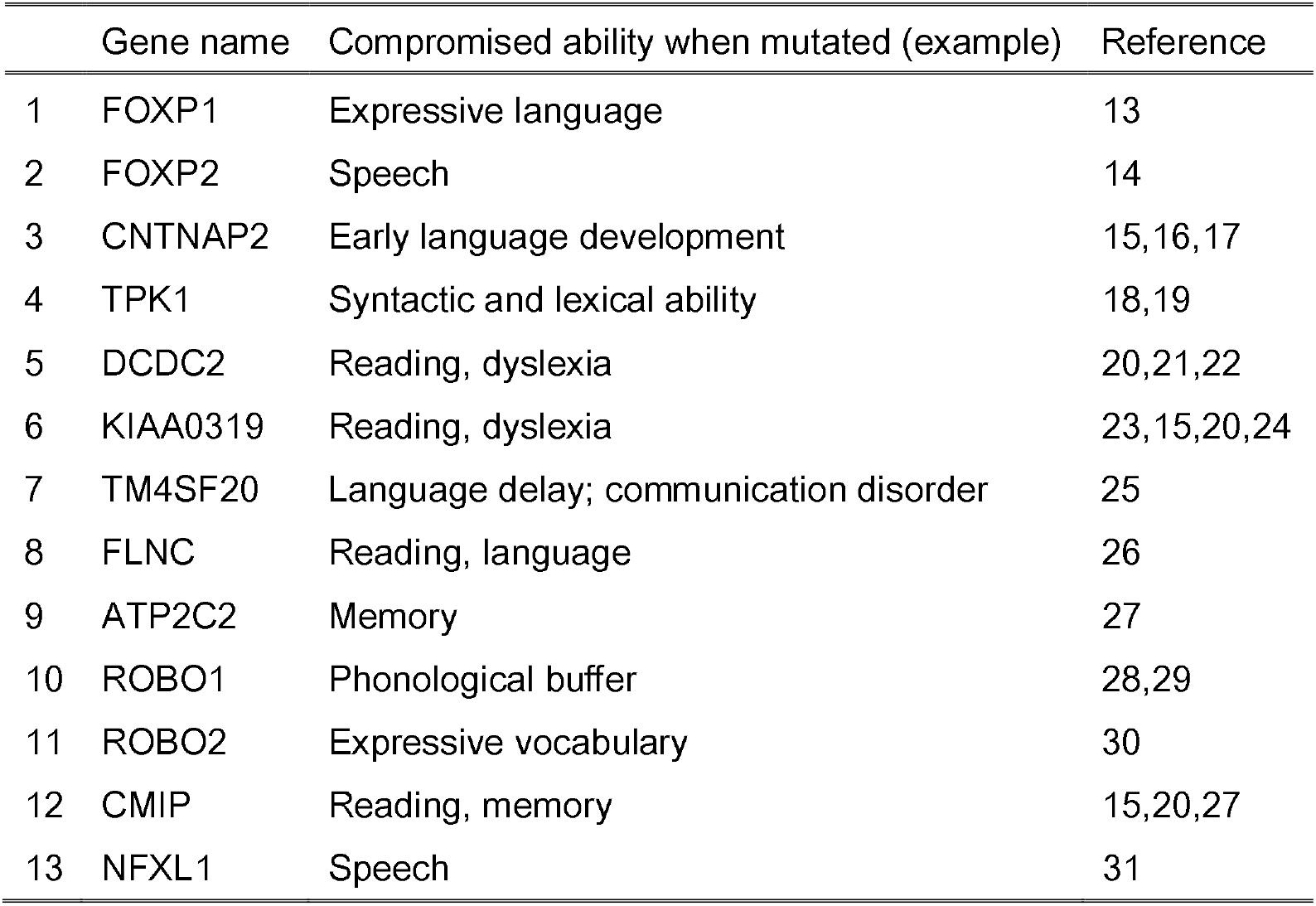
Thirteen language genes

|  | Gene name | Compromised ability when mutated (example) | Reference |
| --- | --- | --- | --- |
| 1 | FOXP1 | Expressive language | 13 |
| 2 | FOXP2 | Speech | 14 |
| 3 | CNTNAP2 | Early language development | 15,16,17 |
| 4 | TPK1 | Syntactic and lexical ability | 18,19 |
| 5 | DCDC2 | Reading, dyslexia | 20,21,22 |
| 6 | KIAA0319 | Reading, dyslexia | 23,15,20,24 |
| 7 | TM4SF20 | Language delay; communication disorder | 25 |
| 8 | FLNC | Reading, language | 26 |
| 9 | ATP2C2 | Memory | 27 |
| 10 | ROBO1 | Phonological buffer | 28,29 |
| 11 | ROBO2 | Expressive vocabulary | 30 |
| 12 | CMIP | Reading, memory | 15,20,27 |
| 13 | NFXL1 | Speech | 31 |

**Table 2.**
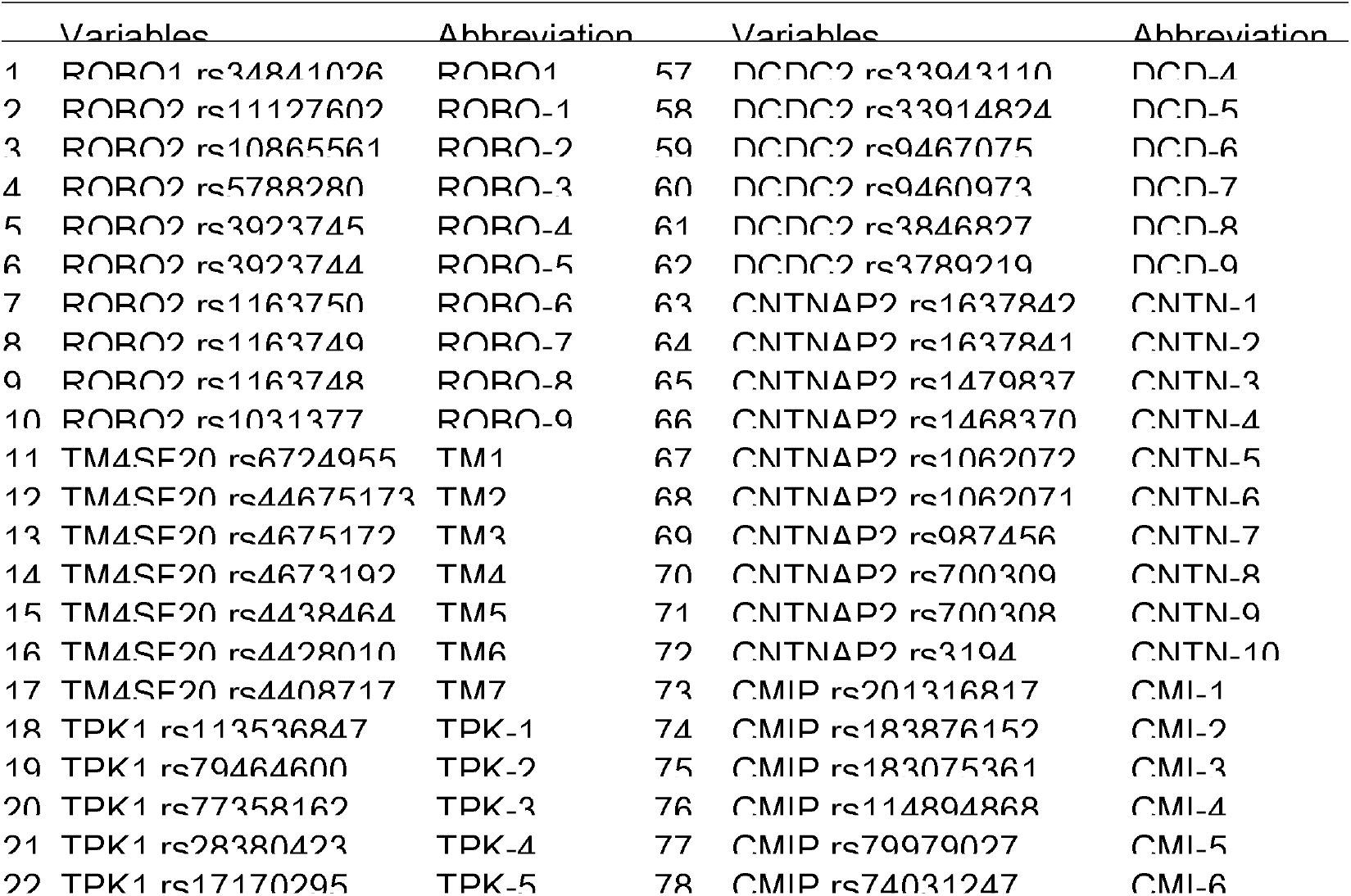
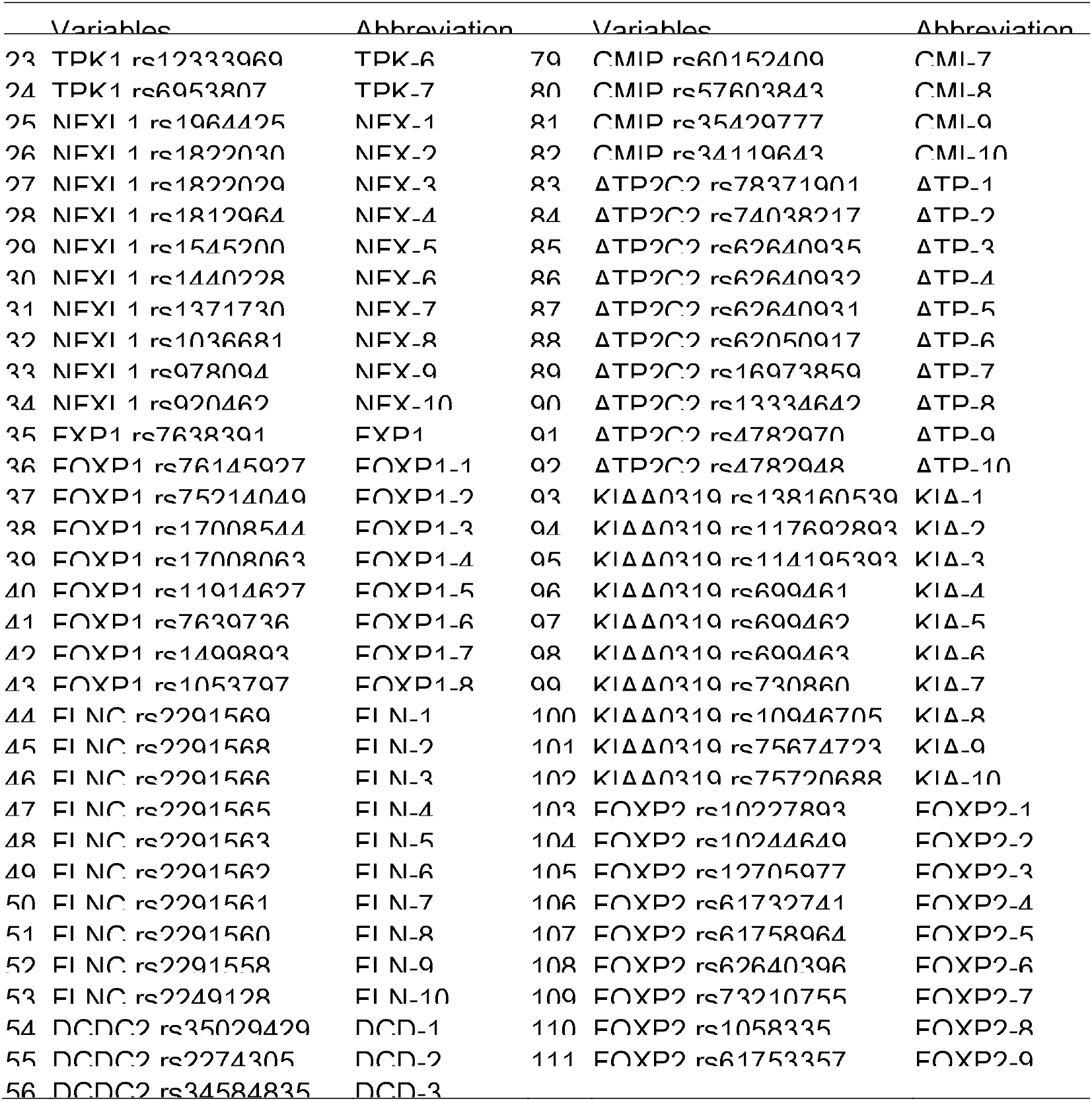
111 SNPs of thirteen language genes

### Collection of culture/ducation parameters

Most education/culture parameters were collected from three websites, including Baidu (https://baike.baidu.com), Bing (https://cn.bing.com), omniglot (https://omniglot.com/writing/languages.htm) and United Nations databases (UND) (http://data.un.org/Default.aspx). (Supplementary file-3) Detailed data are not provided in this manuscript because of page limitation, but can be requested from the corresponding author. This study is a part of our larger scale one that has to collect much more parameters (language gene single nucleotide polymorphism or SNPs, geographical factors, societal factors, etc.) except for education/culture parameters. Besides 29 education/culture parameters (Table 3), we are collecting 111 (SNPs) + 61 (Education/ Culture/ Geography etc.) = 172 parameters in total for around 150 countries, and only 26 countries in Table 4 have been collected for all 172 parameters by the time this manuscript is written.

**Table 3.** Education/culture parameters

| Variables (in number) | Abbreviation | Data source |
| --- | --- | --- |
| 1 Dialect types in the sample country | dial | Baidu, Bing |
| 2 Alphabet of the country's main language | alph | Baidu, Bing, omniglot |
| 3 Vowels of the country's main language | vowe | Baidu, Bing, omniglot |
| 4 Consonants of the country's main language | cnso | Baidu, Bing, omniglot |
| 5 Technology literature | tech | PubMed |
| 6 Engineering literature | engi | PubMed |
| 7 College/institute | inst | Baidu, Bing |
| 8 News agency | news | Baidu, Bing |
| 9 Newspaper/periodicals | newp | Baidu, Bing |
| 10 Broadcast language types | brod | Baidu, Bing |
| 11 Teachers in pre-primary education, both sexes | tpp | UND |
| 12 Teachers in primary education, both sexes | tpe | UND |
| 13 Teachers in secondary education, both sexes | tse | UND |
| 14 Exports of cultural goods | ecgd | UND |
| 15 Imports of cultural goods | icgd | UND |
| 16 Exports of books and press goods | ebpd | UND |
| 17 Exports of performance and celebration goods | epcd | UND |
| 18 Exports of visual arts and crafts goods | evad | UND |
| 19 Natural relics | nrel | UND |
| 20 Cultural relics | crel | UND |
| 21 Natural/cultural double relics | ncrl | UND |
| 22 Total world cultural relics | twcr | UND |
| 23 Power distance Ref. | PDI | Ref,[32-34] |
| 24 Uncertainty avoidance | UAI | Ref,[32-34] |
| 25 Individualism vs collectivism | InCo | Ref,[32-34] |
| 26 Masculinity and Femininity | Masc | Ref,[32-34] |
| 27 Long and short term orientation | Long | Ref,[32-34] |
| 28 Indulgence and restraint | Rest | Ref,[32-34] |
| 29 Cultural context | cuco | Ref,[35] |

**Table 4.** Estimated curo level (ECL) values of selected 26 countries in this study

| Nation/People/Region | ECL | Nations used in this study |
| --- | --- | --- |
| Japanese | 42 | Japan |
| Korea | 40 |  |
| Chinese | 38 | China, Mongolia |
| Vietnam | 36 | Vietnam, Nepal, Myanmar, Bhutan, Sri Lanka, Bangladesh |
| Arab, Middle East, West Asia | 34 | Iran |
| Indian | 32 |  |
| Greek, South European | 30 | Slovenia, Greece, Bosnia and Herzegovina |
| Africa | 28 | Benin, Cameroon, Gambia, Kenya, Senegal, Tunisia, Tanzania |
| Mexican, Middle America | 26 | Barbados, Dominican, Peru |
| Spanish | 24 |  |
| Italian | 22 |  |
| Russian | 20 |  |
| French | 18 | France |
| French Canadian | 16 |  |
| English | 14 | UK |
| English Canadian | 12 |  |
| Australian | 10 |  |
| North American | 8 |  |
| Scandinavian | 6 |  |
| Swiss | 4 |  |
| German | 2 | Germany |

### Selected countries with decent representation

The countries were selected based on a range of cultural context vale (Table 4). In order to give cultural context parameter quantitative estimation (thus can be conveniently used in statistical analysis), the relative cultural context levels for known countries/areas in Table 3 are estimated from 2 to 42 as shown in the column 2. Then other countries were provided similar or same estimated curo level (ECL) value by comparison^[36–38]^. For example, seven African countries (Benin, Cameroon, Gambia, Kenya, Senegal, Tunisia, and Tanzania) were given the same ECL value as 28. Nepal, Myanmar, Bhutan, Sri Lanka, Bangladesh were given the same ECL value as Vietnam because they all belong to south-western Asia.

### Correlation analysis

The software Origin was employed to analyze the potential correlation relationship between all parameters in this study. By using Origin2019, principal components analysis (PCA) was performed to extract potential correlation patterns among multiple numerical variables. PCA is a feasible technique to emphasize variation and visualize strong patterns in a large dataset. In a typical PCA diagram, correlated variables are drawn as short or long arrows in which long arrow represents strong correlation and short arrow represents weak correlation; plus, two arrows can form a right, obtuse or acute angle, representing no-correlation, negative correlation or positive correlation, respectively. Quantitative correlation between any two factors was undertaken as follow. The basic variables to measure in a PCA result plot are arrow length and the angle between two arrows. The correlation score = angle score ×arrow score; the angle score ranges from −5 to +5. The angles (0-15], (15-30], (30-45], (45-60], (60-75], (75-105], (105-120], (120-135], (135-150], (150-165], (165-180] are scored 5, 4, 3, 2,1, 0, −1, −2, −3, −4, −5, respectively. The arrow score = the length of arrow-1× the length of arrow-2. The angle value and the arrow length value can be conveniently obtained with ImageJ software (Supplementary file-4, Supplementary file-5).

## Results and Discussion

### General correlations among all selected SNPs

Figure 1(A) demonstrated a general landscape of correlational relationship among 111 SNPs. Here are several apparent findings. First, no all SNPs from a single gene stay aggregated in a corner of the PCA map, as expected; because all genes passed through million years of mutual adaptation; if all SNPs from a single gene stay aggregated, that would means that many other SNPs from other genes counteract with them. Such a single gene would likely be lost during evolution. Second, we still can find that a couple of (not all) SNPs from a single gene can aggregate at a specific position of the PCA map. Such examples include FLN-2~FLN-3~FLN-5, FOXP1-6~FOXP1-5~FOXP1-7, FOXP1-1~FOXP1-4, ATP-5~ATP-10, CNTN-1~CNTN-6, etc. Third, there are about four big clusters of SNPs, any one of them having a totally negative correlational group and two other groups with much less level of correlations.

**Figure 1.**
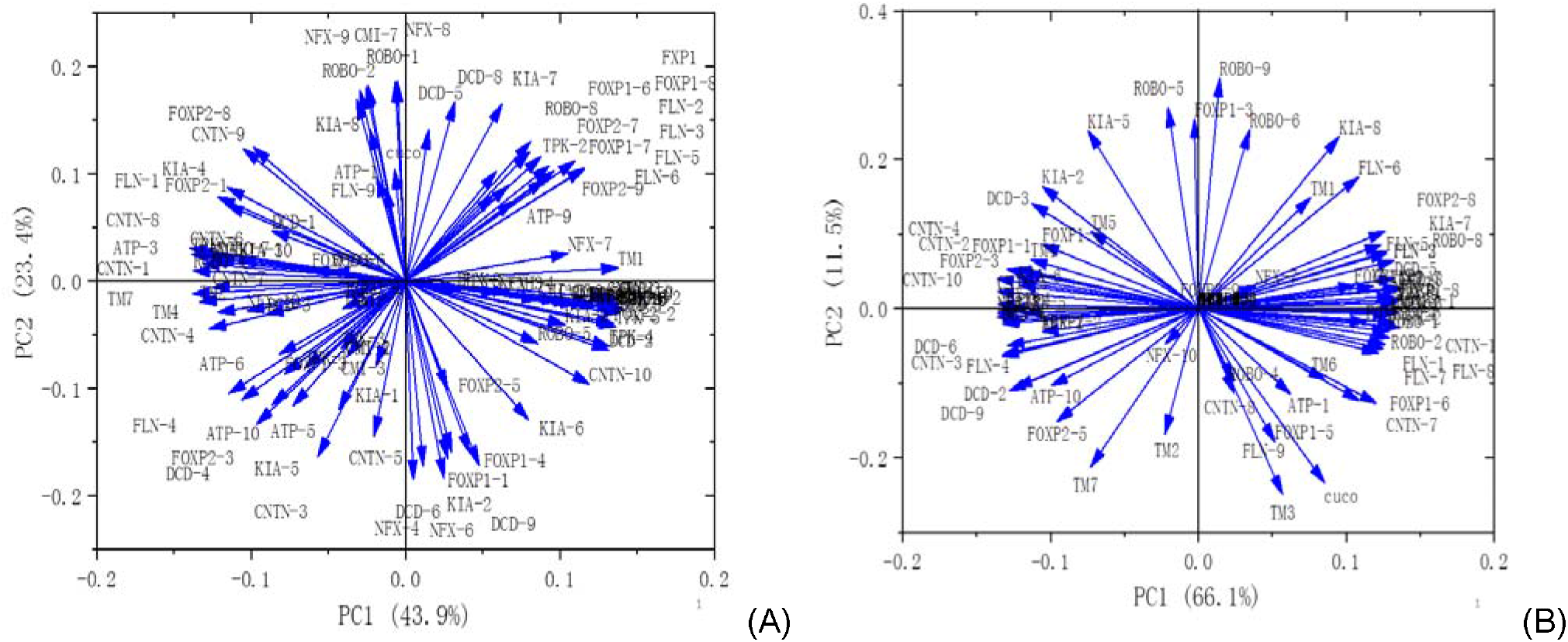
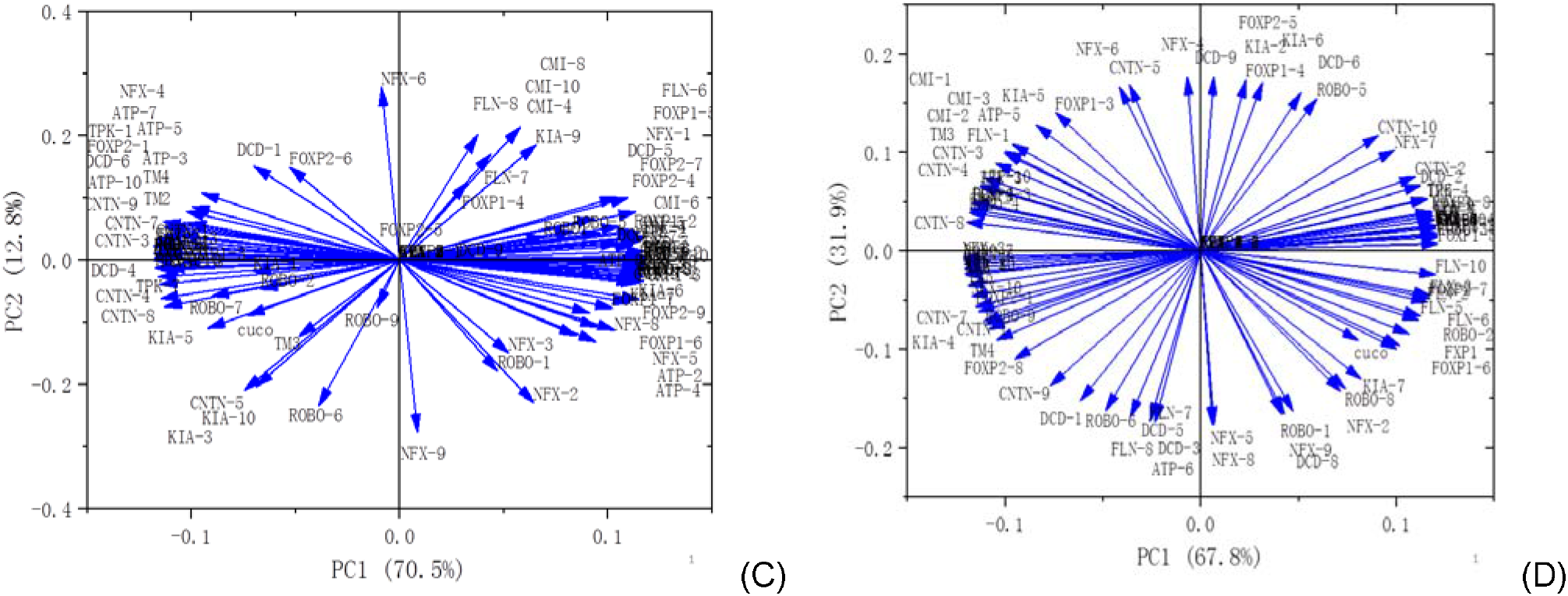
PCA for 111 language gene SNPs. (A) All SNPs in all 26 countries; (B) All SNPs in 10 high cultural context countries; (C) All SNPs in 11 intermediate cultural context countries; (D) All SNPs in 6 low cultural context countries.

### Effect of cultural context (cuco) on correlational patterns among language gene SNPs

Cultural context is an important parameter that can distinguish individual country/region from each other. Countries can be classified as high, intermediate and low cultural context types. In table 4, countries were divided as three groups by cuco values: [2,26], [28,36), and ≥36. Correlation analysis among 111 SNPs was performed for three groups of country, and it was found that SNP correlation pattern changed significantly. In the low cuco group, correlations among language gene SNPs became evenly distributed (Fig.1D), which is not found in high (Fig.1B) and intermediate (Fig.1C) cuco groups.

Table 5 provided parameters that have strongest correlations with cuco in three groups of country. The SNP FOXP1-5 is positively correlated with cuco under high cultural context, but became negatively correlated under intermediate cultural context. The other three SNPs, KIA-5, FLN-6 and CNTN-5, demonstrated similar conversions. Though very limited number of countries was employed for PCA analysis, the above results were still very intriguing, in that this study may provide a good case in which a culture/education factor highly influences genotype interaction pattern(s).

**Table 5.**
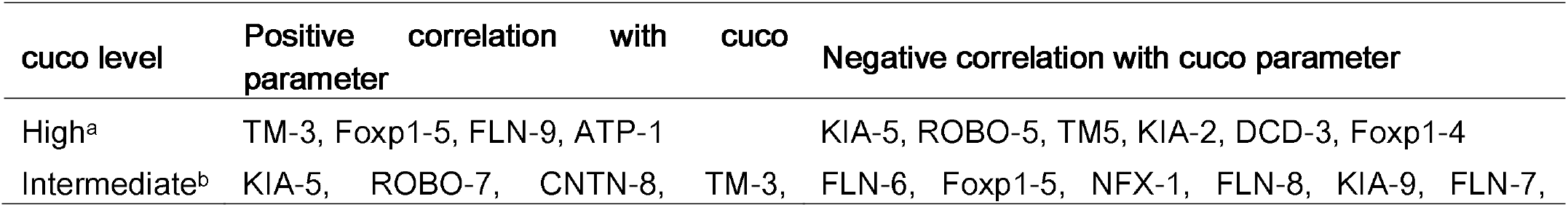
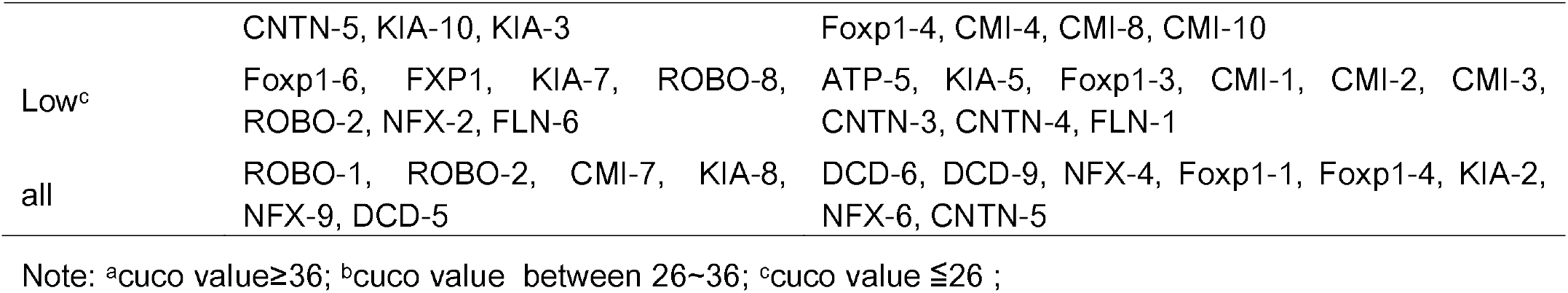
Parameters that have strongest correlations with cuco in three groups of country

### Correlation between a specific gene SNP and culture/education parameters

Correlation analysis among all variables in Table 2 and Table 3 is of interest. In theory, culture/educational activities still belong to human phenotypes, while any phenotype will automatically bring feedback to genotypes that determine it. So the long-run human culture/education factors shall somehow interact with gene polymorphisms and form related (weak or strong) functional correlations.

In Figure 2(A), strongest positive correlations were seen at ATP-1-tech, ATP-1-ecgd, ATP-1-engi, ATP-1-tpp and ATP-1-evad. ATP-1 is one of the SNPs of language gene ATP2C2, which encodes the ATPase secretory pathway Ca2+ transporting-2 protein. Diseases associated with ATP2C2 include specific language impairment and speech/ communication disorders. The five parameters, tech (Technology literature), engi (Engineering literature), ecgd (Exports of cultural goods), tpp (Teachers in pre-primary education, both sexes) and evad (Exports of visual arts and crafts goods), have similar highest correlation values. It is surprising that the five strongest correlations were all formed by ATP-1 SNP. How these five parameters mechanically correlate with the language gene SNP ATP-1 will be worth tackling in the future.

**Figure 2.**
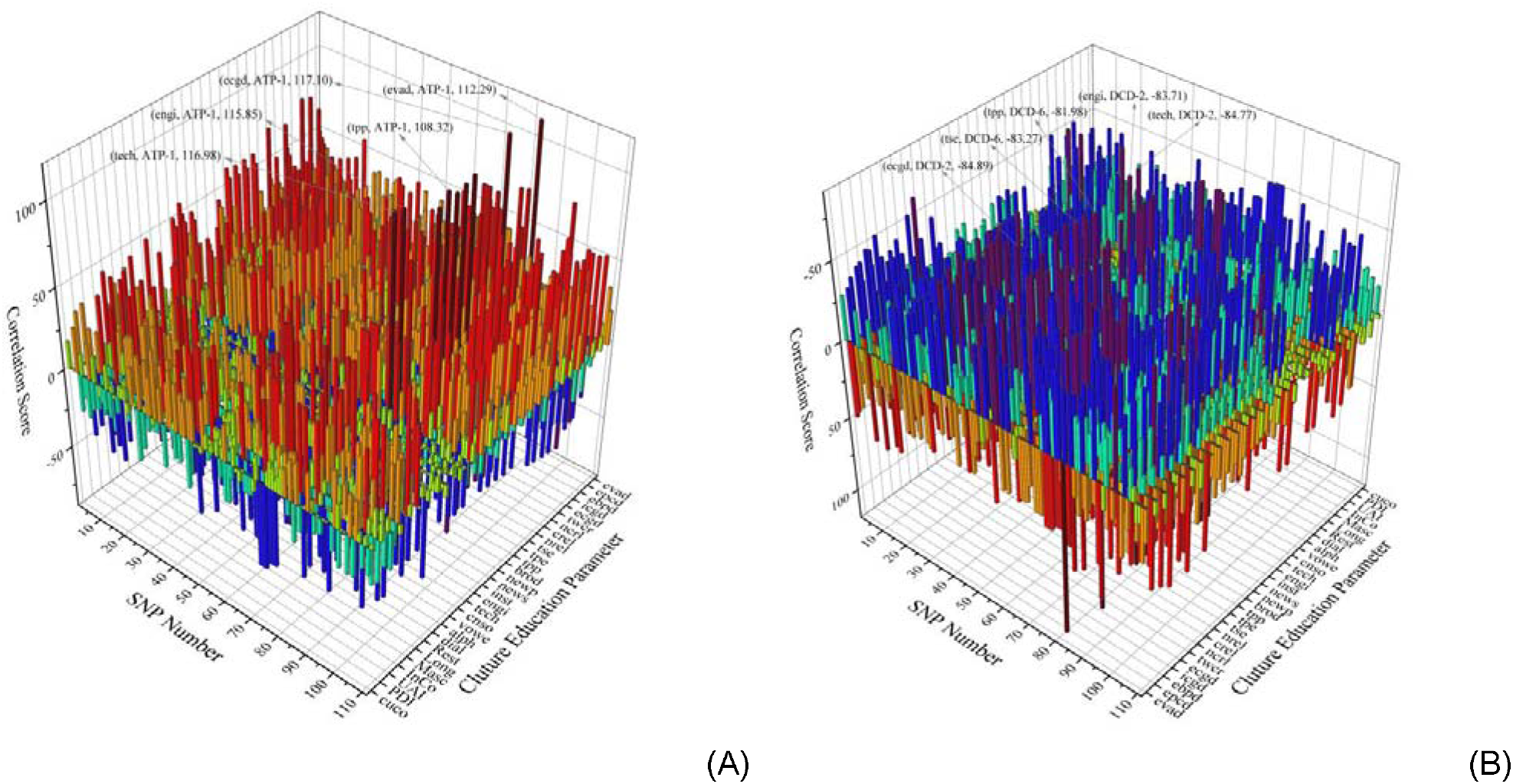
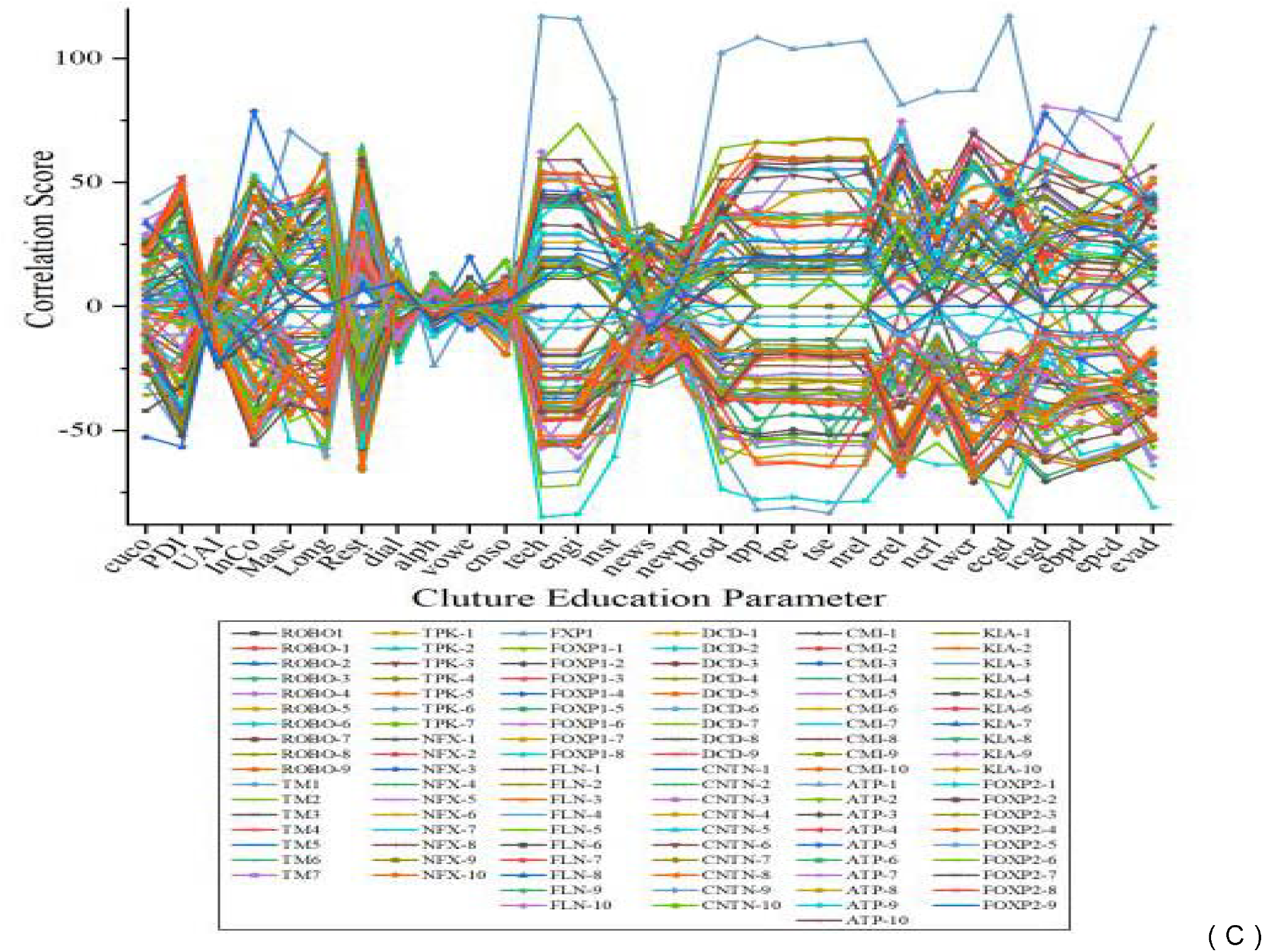
Correlation among language gene SNPs and 30 culture/education parameters. (A) Five strongest positive correlations were shown; (B) Five strongest negative correlations were shown; (C) Smallest correlations were exposed clearly at five culture/education parameters.

In Figure 2(B), strongest negative correlation were seen at DCD-2-tech, DCD-2-engi, DCD-6-tse (Teachers in secondary education, both sexes), DCD-2-ecgd, and DCD-2-evad. DCD-2 and DCD-6 are two SNPs of language gene DCDC2.This gene encodes a doublecortin domain-containing family member, which is thought to function in neuronal migration where it may affect the signaling of primary cilia. Mutations in this gene are associated with reading disability (RD) type 2 (developmental dyslexia)^[40]^. Robert Plomin^[39]^ claimed that a person’s achievement in English, mathematics, science, literacy and arts is significantly heritable and influenced by similar groups of genes. They compared 12,500 pairs of twins’ genome sequences and exam available scores of all school tests. One SNP locus (rs807701, not the same DCDC2 SNP as in this study) on DCDC2 gene seemed highly associated with reading ability, and previous studies already found that DCDC2 mutations will lead to reading difficulty^[40]^. The large scale sampling of the above study enabled the authors to abstract relatively confident conclusions on how genes affect a person’s academic performance, including language abilities and general output in the liberal education^[41]^. Such performance is supposed to have a good chance to influence people’s behavior in cultural/educational activities.

Figure 2(C) indicates that little correlation was found for UAI (Uncertainty avoidance), dial (Dialect types in the sample country), vome (Vowels of the country’s main language), alph (Alphabet of the country’s main language) and cnso (Consonants of the country’s main language); According to Hofstede^[32–34]^, uncertainty avoidance describes the extent to which people attempt to cope with anxiety by minimizing uncertainty. Cultures with high score in uncertainty avoidance prefer rules and structured disciplines, and employees tend to keep longer with their current employer. UAI may be a universal parameter in any country/region though its content differs a lot in different country/region, and that may be why it has a very small correlation value with all SNPs; The other four parameters, dial, vome, alph and cnso, are all basic language factors. It is interesting why they also have very small correlation values with all language gene SNPs.

### Conclusions

In the past several years the authors have been collecting multiple parameters (history /geography/religion/ genetics/ education/ culture/ society) for many countries, and started to supply with human language gene polymorphism data plus appropriate correlation analysis^[38,42–43]^. Such multi-layer data, when in a scale large enough, shall be useful for investigations on interdisciplinary questions on the boundary of natural science and social science. Recently, researchers found interesting interplay of genetics and culture in Ethiopia^[7]^, highlighting the importance to employ data from both natural science and social science to deepen the understanding of cultural questions.

In this study, the basic data include 13 language genes and their randomly selected 111 single nucleotide polymorphisms (SNPs), SNP profiles in 26 countries, 29 culture/education parameters in 26 countries, and estimated cultural context values for 26 countries. In order to undertake principal component analysis (PCA), SNP genotypes, cultural context and all other culture/education parameters have to be quantitatively represented into numerical values.

In this study, only 26 countries were used for all analysis. However, the namelist (Table 4) contains descent representation, including one eastern (Asia) developed country, three western developed countries, two developing countries in eastern Asia, three ordinary European countries, seven African countries, six ordinary countries in south Asia, and several typical ones in west Asia (one) and Middle America (three). So the results in this study would be preliminary but valuable.

Based on the above conditions, this study obtained its preliminary results, the main points of which contain: (1) The 111 SNPs form several clusters of correlational groups with positive and negative correlations with each other; (2) Cultural context level apparently influences the correlational patterns among 111 SNPs in the principal component analysis diagram; and (3) Among 29 culture/education parameters, several basic characteristics of a language (the numbers of alphabet, vowel, consonant and dialect) surprisingly demonstrate least correlations with 111 SNPs of 13 language genes.

## ACKNOWLEDGMENTS

This study was supported by National Research Center for Foreign Language Education Grant (ZGWYJYJJ10A042) and State Language Commission Research Grant (YB135-117). The authors thank Nie Qi, Pan Yusheng for their help in collecting sample data.

## Supplementary files

Supplementary file-1 Selected 111 SNPs from 13 language genes

Supplementary file-2 SNP quantification method and SNPs data in 26 countries

Supplementary file-3 Culture education parameters data

Supplementary file-4 PCA for language gene and culture education parameters

Supplementary file-5 SNP and culture education correlation value data

